# Massively parallel functional screen identifies thousands of regulatory differences in human versus chimpanzee postcranial skeletal development

**DOI:** 10.1101/2025.10.21.683789

**Authors:** Alexander S Okamoto, Clarissa R Coveney, Danalaxshmi S Ganapathee, Terence D Capellini

## Abstract

Every element of the human skeleton exhibits some differences in comparison to our closest living relatives, chimpanzees. Many of these skeletal modifications underpin key events in human evolution, enabling our species to walk upright, manipulate tools with precision, and support enlarged brains. Identifying the genomic changes that underlie these features remains an outstanding challenge due to the substantial number of differences between the human and chimpanzee genomes. To identify human-chimp sequence differences that modulate gene expression in the developing postcranial skeleton, we used a massively parallel reporter assay (MPRA) to screen the human and chimp versions of 70,000 regulatory elements present in the prenatal skeletal template for differential activity. After testing our library in two cartilage and one bone marrow-derived lymphoblast line, we identify 30,736 regions (45.2%) with activity in our assay. Of the active regions, we find that 11,542 (37.6%; or 17% of the entire pool) regions exhibited differential activity between the human and chimpanzee. We find that human ancestor quickly evolved regions (HAQERs) were predictive of differential activity while Human Accelerated Regions were not and both sets failed to predict the magnitude of effect, unlike the total number of base pair differences between species, which was weakly correlated with effect size. These findings reveal that human skeletal evolution involves widespread regulatory changes distributed across thousands of elements rather than concentrated effects at a few key loci, supporting a polygenic model for the evolution of complex morphological traits.

## Introduction

The human postcranial skeleton exhibits numerous adaptations for derived traits including bipedal locomotion, extensive tool use, and extended growth periods (Darwin 1871). Indeed, nearly every element of the human skeleton exhibits some species-specific features in comparison to our closest living relatives, chimpanzees (Pan troglodytes). While some of these changes are due to neutral processes such as genetic drift, many are likely adaptations for brain, locomotion, diet, and life history changes during hominin evolution. Understanding which genomic differences between humans and chimpanzees actually drive these morphological distinctions represents a fundamental challenge in evolutionary biology.

The task of identifying the genomic basis of human-specific features is complicated by the substantial number of differences between the human and chimpanzee (*Pan troglodytes*) genomes (Varki and Altheide 2005). Despite sharing 99% sequence identity when comparing single-nucleotide variants (SNVs), humans and chimpanzees differ at approximately 30 million nucleotide positions in a haploid genome consisting of 3 billion base pairs (Mikkelsen et al. 2005; Yoo et al. 2025). A further 90 million base pairs differ between the genomes due to structural variation (indels), accounting for another 3% of the total genome (Mikkelsen et al. 2005). Most of these differences fall within non-coding regions of the genome, suggesting that they cause differences in gene expression between the two species due to alternations in the activity of regulatory elements (King and Wilson 1975). By comparing the regulatory activity of orthologous genomic sequences between these species, we can identify which specific DNA changes may have contributed to the developmental programs underlying human skeletal evolution.

A powerful tool for identifying human-specific sequences impacting gene expression via *cis*-regulation is the massively parallel reporter assay (MPRA) (Gallego Romero and Lea 2023; Rong et al. 2024; Easterlin and Ahituv 2025). MPRAs allow for the simultaneous screening of many thousands of sequences for regulatory activity as well as differential regulatory activity between paired sequences, such as orthologous regulatory element sequences from human and chimpanzee. Accordingly, studies seeking to identify genomic regions based on sequence features that are likely to underlie modern human traits – such as an excess of human-specific fixed substitutions, losses of conserved sequence, or ancient introgressed variants – have increasingly relied on MPRAs to validate identified sequences and examine their functional effects (Weiss et al. 2021; Jagoda et al. 2022; Whalen et al. 2023; Xue et al. 2023). MPRA results largely depend on the regulatory environment of the tested cell-type (Siraj et al. 2024), requiring a close match between the natural regulatory context of tested sequences and the cell-type used for transfection. When testing human-specific sequences identified from genomic comparisons, neuronal cell lines are most frequently used due to the extent of brain evolution in the hominin lineage (Uebbing et al. 2021; Girskis et al. 2021; Mangan et al. 2022; Pizzollo et al. 2022). While such studies have revealed important aspects of human-specific *cis*-regulatory changes involved in neurodevelopment, comparing the relative efficacy of various alternate methods for determining human-specific genomic features across studies is challenging due to differences in the MPRA protocols used in each study. This leads to an incomplete picture of how effectively different genomic features predict gene expression differences. Comprehensive studies (i.e., testing all types of genetic changes in the same assay) have not been conducted in any cell type, hampering our ability to understand the differing impacts that qualitatively different types of genetic modifications have on gene expression and phenotypic biology.

Therefore, to address the above limitations, we focused on the regulation of skeletal development. Skeletal cell types are dramatically less diverse than neurological cell types and therefore provide an opportunity to comprehensively test human/chimp sequence changes in fewer cell types relevant to skeletal morphology. We focused in particular on the postcranial skeleton because it forms primarily endochondrally, with many human-specific features first observed at the cartilage level during gestation (de Bakker et al. 2016; Young et al. 2022; Senevirathne et al. 2025). We integrated datasets on chromatin accessibility in chondrocytes (Richard et al. 2020; Young et al. 2022; Richard et al. 2025; Okamoto et al. 2025) sampled from across the developing human postcranial skeleton to comprehensively screen skeletal regulatory elements for differential activity between humans and chimpanzees via MPRA (Figure 1). As the cartilage model prefigures adult bone its varied skeletal cells consist of growth plate chondrocytes localized to the shaft of the element, and epiphyseal and articular chondrocytes along the element ends. We therefore test our comprehensive set of developmental human skeletal regulatory elements in three cell lines, two chondrocyte (one growth plate and one epiphyseal/articular) and a bone marrow-derived lymphoblast line. In each cell line, we identify regulatory elements for which the human and chimpanzee sequences drive differential enhancer activity and test the relative efficacy of multiple methods of identifying human-specific sequence features for predicting human-specific enhancer activity across the developing skeleton. This dataset allows us to directly compare alternative regulatory element sets overlapping different human-specific features, such as human accelerated regions (HARs) or human ancestor quickly evolved regions (HAQERs) for differential activity between human and chimp (Pollard et al. 2006; Prabhakar et al. 2006; Bird et al. 2007; Bush and Lahn 2008; Gittelman et al. 2015; Kostka et al. 2018; Mangan et al. 2022; Bi et al. 2023; Whalen et al. 2023; Keough et al. 2023; Yoo et al. 2025).

**Figure 1.**
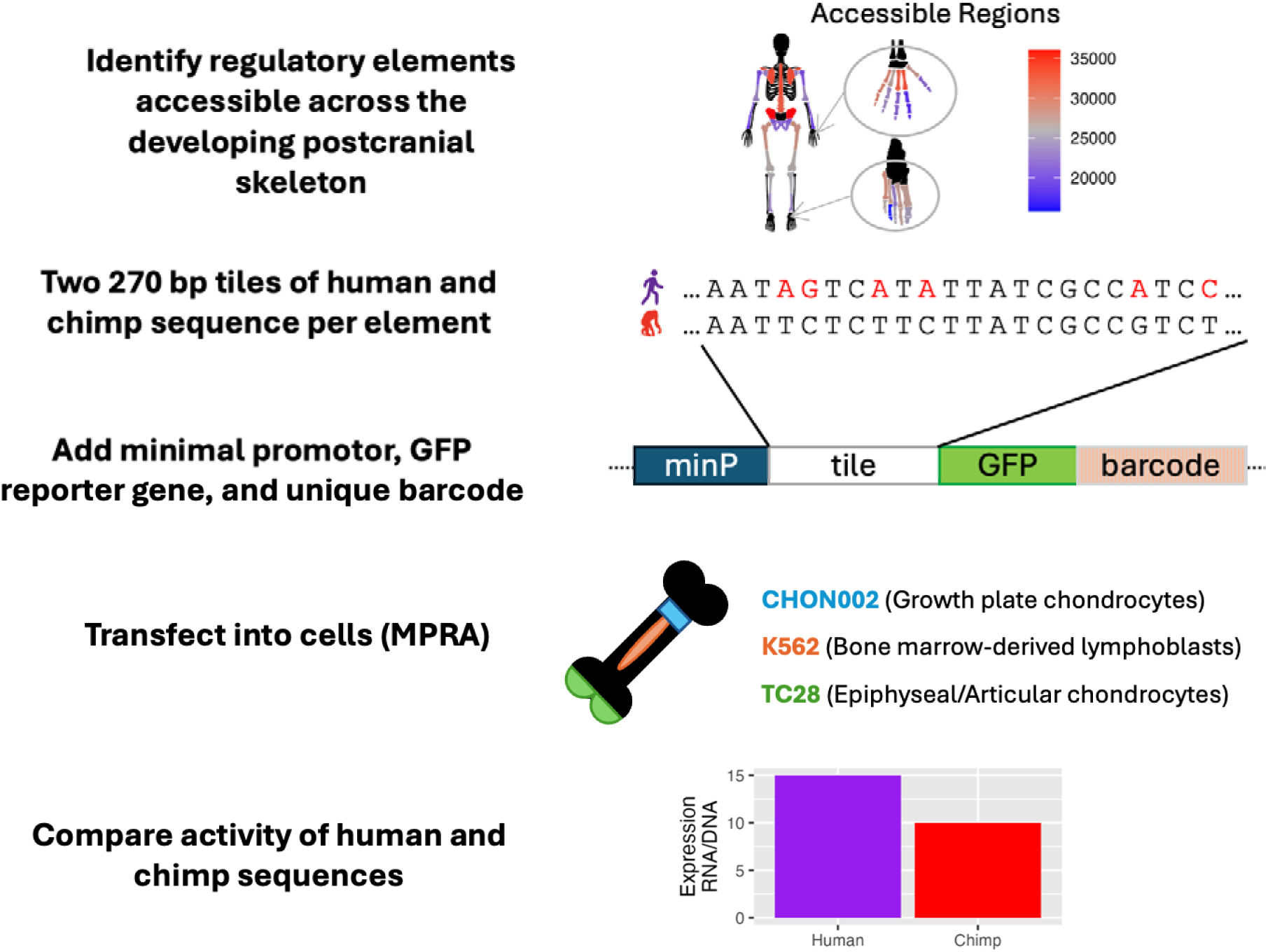
MPRA experimental design. Regulatory elements accessible across the developing human skeleton were standardized to a length of 540 base pairs and divided into two tiles to design an MPRA library that included both the human and chimpanzee sequences for each tile. This library was then transfected into three cell lines to measure the relative potential of the human and chimp sequences to drive reporter gene expression (*GFP* – green fluorescent protein).

## Results

We previously sampled 37 distinct postcranial regions from the developing human skeleton between gestational days (E) 50-70, a window of time in which the individual elements of the skeleton undergo rapid cartilage morphogenesis and establish the basic architectural framework that will define adult skeletal morphology (Richard et al. 2020; Young et al. 2022; Richard et al. 2025; Okamoto et al. 2025). Using the assay for transposase accessible chromatin followed by next generation sequencing (ATAC-seq) on chondrocytes from these regions, we identified 80,460 genomic regions that are accessible in at least one skeletal element and not in embryonic brain. This brain filter is intended to remove regulatory elements likely controlling cellular housekeeping processes common to all cell types and in our experience filtering by additional distinct tissue datasets does not markedly reduce these signals (Young et al. 2022). After removing ENCODE blacklisted regions (Amemiya et al. 2019), and those that were identical in sequence between humans and chimpanzees, we generated a set of remaining skeletal cell ATAC-seq regions that displayed any sequence difference between humans and chimpanzees. These 77,084 regions were then used to generate an MPRA oligonucleotide library, in which we created two matched, 270 base pair tiles (one chimpanzee and one human sequence oligo) for each region (144,815 total tiles). We then tested these 289,630 sequences in the MPRA in two distinct skeletal cell types - T/C-28a2 and CHON002 chondrocytes, with cellular properties better reflecting articular chondrocyte biology and growth plate chondrocyte biology, respectively – and one non-skeletal bone-marrow derived cell type – K562 cells, an immunological precursor (TC28, CHON002, and K562, henceforth; Figure 1).

We performed five replicates of library transfection in each cell type. Overall, counts across replicates correlated highly (Pearson’s R≥0.84) except for one replicate of K562, which was excluded from further analyses (Figure S1). We found that our positive controls were highly active in K562 (89%) and TC28 (77%) but less active in CHON002 (45%) (Table S1). This is expected as the controls used in this study have been previously validated in K562s and other lymphoblastoid cell lines rather than chondrocytes (Tewhey et al. 2016; Siraj et al. 2024).

Negative controls showed low rates of activity (K562: 8%, TC28: 1%, CHON002: 0%), confirming that our activity thresholds successfully distinguished active from inactive sequences.

### Nearly half of skeletal regulatory elements show functional activity

After excluding poorly aligned sequence pairs, we analyzed a total of 117,031 paired tiles falling within 67,986 genomic regions accessible in the developing skeleton (hereafter, regions; Figure S2). Of these regions, 30,736 (45%) had activity in at least one tile for one or more cell-types and are likely enhancers that upregulate target genes (Tewhey et al. 2016) (Figure 2A). See Table S2 for tile statistics. Overall, the greatest number of active regions was identified in K562 cells, followed by TC28 and then CHON002. Activity differed substantially between cell types, with only 2,162 regions (7%) active in all three cell types, though many regions were active in both K562 and TC28 cells (34% of active regions, Figure 2B). Some regions showed activity only in one cell type (993 uniquely active in CHON002, 2,327 in TC28, 10,414 in K562). Considering all sequences together, no cell type showed a bias towards activity in human or chimp tiles (P > 0.05, two-tailed binomial test).

**Figure 2.**
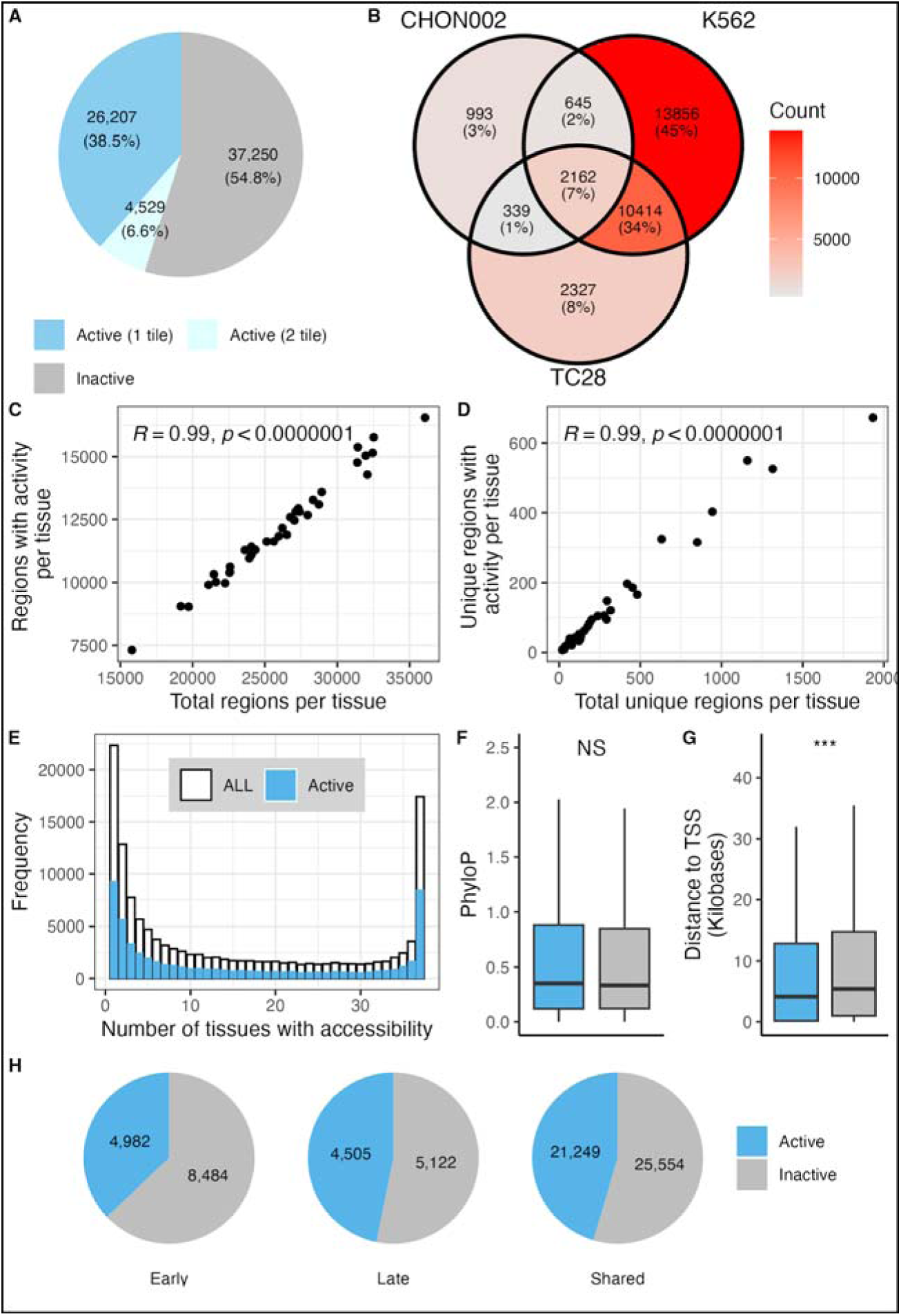
Characteristics of active regions. (A) pie chart of the number of tested regions containing one or two active tiles. (B) Venn diagram showing the number of tested regions sharing activity across cell types. (C) correlation of elements with activity in any cell type per tissue with the total number of tested regulatory elements for that tissue. (D) correlation of tissue-unique elements with activity in any cell type with the total number of tested unique regulatory elements for that tissue. (E) frequency of all test regions and active elements falling in regions with different levels of shared accessibility. (F) phylogenetic conservation (PhyloP scores) of active versus inactive tiles, NS = not significant. (G) distance to the nearest transcription start site (TSS) for active versus inactive regions. *** = P < 2.2 x 10-16, Wilcoxon rank-sum test. (H) pie chart showing activity for regions accessible at different developmental stages (early, late, or both early and late stages).

Of the active elements, both tiles showed activity for 4,529 regions (15%) compared to 26,207 regions with activity in only a single tile (85%; Figure 2A). Out of those elements with activity across both tiles, 3,833 were identified within the same cell line whereas in 696 regions, the two tiles were active in different cell types. For the regions with activity across both tiles in a single type, activity level was significantly correlated for both human and chimp sequences (Pearson’s correlation, P < 0.05 for each cell line), except chimp tiles in CHON002 for which the correlation is not significant but trends in the same direction (R = 0.14, P = 0.18, Figure S3A).

By contrast, comparing tiles with activity in different cell types revealed no correlation for either human or chimp, suggesting these regions may contain two distinct enhancer elements, one falling within each adjacent tile (Pearson’s correlation, P > 0.05 for each pairwise comparison of cell lines; Figure S3B).

To explore the biological features of sequences with activity in this assay, we first identified the number of active regions with accessibility in each of the 37 skeletal elements used to generate this dataset. We found a strong correlation between the number of regions with accessibility in each tissue and the subset of those elements with activity in the MPRA (Figure 2C). This pattern holds when analyzing element sets with accessibility uniquely in one tissue as well as across all three cell types (All Pearson R≥0.94, *P< 1.0×10^-7^,* Figures 2D, S4).

As many regulatory elements are accessible in multiple parts of the developing skeleton, thereby driving pleiotropic gene expression, our MPRA results might show a bias towards greater rates of activity for regions with accessibility in more tissues, which are more likely to be accessible in the cell lines used in this experiment. Instead, we found that active elements show a similar distribution of tissues with accessibility to the total tested set, with high rates of tissue-specificity or accessibility in all tissues (Figure 2E). This was true for all three cell types (Figure S5A). This consistency suggests that none of the tested cell lines show a strong bias towards any of the developmental tissues used to design this library and may reflect the episomal nature of this assay (see Discussion).

We then examined the role of evolutionary sequence conservation on activity patterns. We found that active tiles did not show higher levels of sequence conservation than inactive tiles (Wilcoxon rank-sum test, P = 0.14, Figure 2F) as measured by average phyloP scores per tile. Therefore, recently evolved human sequences were just as likely to show enhancer activity as mammalian conserved regions in this experiment, as is expected given that the regulatory elements were chosen due to their accessibility in human tissues and tested in human cell lines. Regions with activity were generally located closer to transcriptional start sites than inactive regions, consistent with the idea that active regions act as enhancer elements to nearby genes (Wilcoxon rank-sum test, P < 2.2 x10-16, Figure 2G). We also tested for a bias in activity due to the timepoint of accessibility in human development; we partitioned our tested tiles into those with accessibility in at least one tissue at only a single time point (early, ∼E54, 13,466 regions; late, ∼E67, 9,627 regions) or at both timepoints (46,803 regions). Overall, regions with accessibility at the late timepoint or both time points (46.8% & 45.4% active, respectively) had higher rates of activity compared to regions accessible only at the early timepoint (37.0% active), suggesting that the cell lines used in this experiment better capture the environment of later-staged developmental tissues (Figure 2H).

To identify biological pathways regulated by active elements, we performed gene ontology enrichments using the genes linked to elements active in each cell type (McLean et al. 2010). Compared to the total set of oligos, K562 active regions had no gene ontology enrichments, TC28 active regions were enriched for “MHC protein complex”, and “peptide antigen binding,” and CHON002 active regions were enriched for five terms: “cytoplasmic pattern recognition receptor signaling pathway in response to virus”, “CBM complex”, “MOZ/MORF histone acetyltransferase complex”, “granular component”, and “box C/D snoRNP complex” (Tables S3-5), suggesting that activity was not highly biased towards a limited set of biological pathways. Finally, we sought to identify the transcription factors (TFs) driving activity in each cell type. We found that the human sequences of active tiles in CHON002, K562, and TC28 were enriched for 164, 179, and 171 TF binding motifs, respectively, compared to all tested tiles (Table S6). 109 motifs were enriched in all cell types, but some were specific to each cell type, including *SREBF(var.2), ZNF135*, and *MEF2D* in CHON002, *GATA5, GATA::TAL1*, and *GATA2* in K562, and *TP63, RFX2,* and *RFX5* in TC28 (top 3 most enriched unique TF binding motifs). When this analysis was repeated with the chimp active tiles, results were largely similar (Table S7), and no TFs showed differential enrichment between the tiles active in each species for any cell type. Human active sequences were consistently depleted in all three cell lines for *NFIC, NFIA, NFIX* binding motifs compared to the total library, suggesting that one or more these TFs may be important for the activity of the sequences *in vivo* but could not be tested in this experiment due to the transcriptional environment of the cell types used.

### Over one-third of active sequences show differential activity between human and chimp

Of the active regions, 11,542 (37.6%; or 17% of the entire pool) exhibited statistically significant differential activity between the human and chimpanzee versions of at least one tile (Figure 3A). We refer to regions containing at least one differentially active tile hereafter as differentially active regions. This represents a substantial number of functional regulatory differences between the species, suggesting that regulatory evolution has been widespread rather than concentrated in specific genomic regions or pathways. Like the overall activity pattern, differential activity was often distinct to a single cell type. We found 1,180 differentially active regions in CHON002 (28% of active sequences), 9,043 in K562 (31%), and 3,883 in TC28 (24%). 195 regions were differentially active in all three cell types (2%), with 9,137 differentially active in only a single cell type (Figure 3B). As seen with activity, K562 and TC28 cells shared the most differentially active regions. Tiles differentially active in multiple cell types showed a consistent direction of effect in almost all cases (96.6%, Figure S6A). Importantly, differentially active tiles for TC28 and K562 show a bias towards higher expression in human than chimp (P < 0.0001 & P = 0.02, respectively, two-tailed binomial test) while no difference was found in CHON002 (P = 0.5, Figure S6B). 615 regions contained two differentially active tiles (5% of active regions). While most of these regions showed differential activity across both tiles in the same cell type (459 regions), 156 regions contained tiles with differential activity in different cell types (30 TC28-CHON002, 60 TC28-K562, and 104 K562-TC28 regions). The skews between the human and chimp sequences with differential activity across both tiles in the same cell line were tightly correlated in TC28 and K562 (Pearson’s R = 0.36, P < 0.01, R = 0.11, P < 0.05, respectively) but not CHON002 (R = 0.03, P = 0.38) (Figure S6C). Skews were not correlated in tile pairs differentially active in different cell types (Figure S6D).

**Figure 3.**
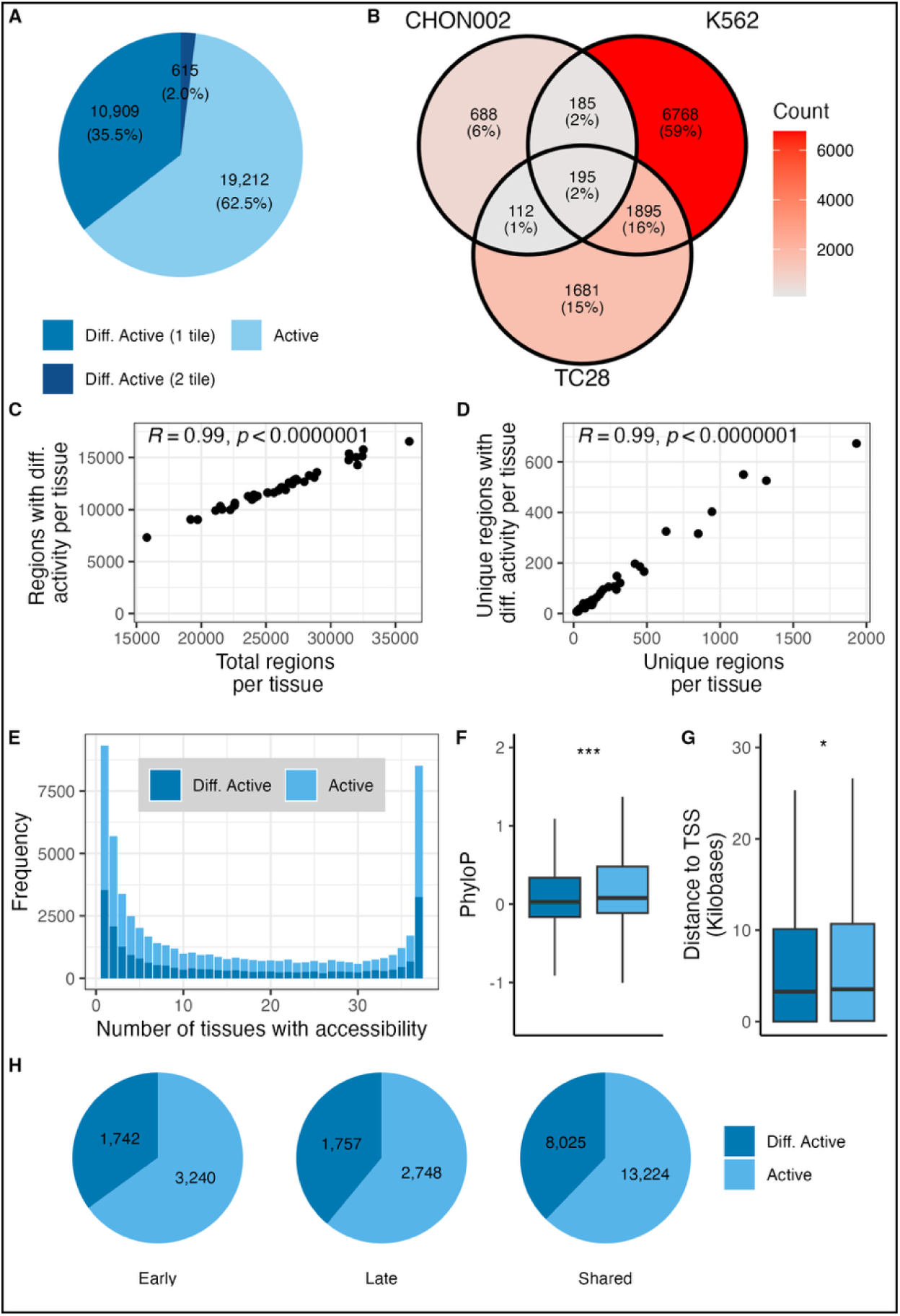
Characteristics of differentially active regions. (A) pie chart of the number of active regions containing one or two differentially active tiles. (B) Venn diagram showing the number of regions sharing differential activity across cell types. (C) correlation of elements with differential activity in any cell type per tissue with the total number of tested regulatory elements for that tissue. (D) correlation of tissue-unique elements with differential activity in any cell type with the total number of tested unique regulatory elements for that tissue. (E) frequency of all active regions and differentially active elements falling in regions with different levels of shared accessibility. (F) phylogenetic conservation (PhyloP scores) of differentially active versus active tiles. (G) distance to the nearest transcription start site (TSS) for differentially active versus active regions. * = P < 0.01, *** = P < 2.2 x 10-16, Wilcoxon rank-sum test. (H) pie chart showing differential activity for active regions accessible at different developmental stages (early, late, or both early and late stages).

The differentially active regions show similar patterns of accessibility across the skeleton to that seen in the active regions, and accordingly, the number of differentially active regions correlates highly with both all regions accessibility in each tissue as well as those uniquely accessible in each tissue (Figure 3C-D). These patterns are consistent across all three cell types (All Pearson R≥0.94, *P< 1.0×10^-7^,* Figures S4B,D, S5). PhyloP scores were significantly lower for active sequences that were differentially active compared to those that were not (Wilcoxon rank-sum test, P < 2.2 x 10-16, Figure 3F). This is likely explained by the fact that higher rates of differential activity are observed in more divergence sequences which are expected to have lower conservation scores (see below). Differentially active regions were also located significantly closer to transcriptional start sites (Wilcoxon rank-sum test, P < 0.01, Figure 3G). Regions active at only one timepoint or across both timepoints had similar rates of being differentially active between human and chimp (35% early, 39% late, 38% shared, Figure 3H).

Against the backdrop of active sequences in each cell type, differentially active sets showed no functional annotation enrichments via GREAT (McLean et al. 2010). When the total set of differentially active regions was tested against a whole genomic background, GREAT performed poorly as 44% of all genes were implicated, although some skeletal pathways were identified (Table S8). When we compared the TF enrichments in the human sequences of differentially active tiles with those of all active tiles, the only enrichment was for *CTCF* in K562 cells (Table S9). No enrichments were identified for the chimpanzee sequences, implying that changes to *CTCF* binding may explain differential activity in some of the tested elements. Direct comparison of all human and all chimpanzee sequences for differentially active tiles identified no significant TF motifs.

### Predictive value of sequence features associated with human evolution

Our comprehensive dataset enabled direct testing of computational approaches for predicting functionally important human-specific sequences. We analyzed 197 regulatory elements overlapping human accelerated regions (HARs) and 61 overlapping human ancestor quickly evolved regions (HAQERs), representing established methods for identifying rapidly evolving genomic segments. Due to the design of this MPRA experiment, another set of human-specific sequences, human conserved element deletions (hCONDELs) (McLean et al. 2011; Xue et al. 2023), could not be investigated (but see Supplementary Text, Figure S7). Out of the 57 active regions overlapping a HAR and 30 active regions overlapping a HAQER, 19 and 19, respectively were differentially active (Figure 4A). One differentially active region overlapped both a HAR and a HAQER (chr8:1700994-1701534). Within all active tiles, those overlapping a HAR were not more likely to be differentially expressed than expected by chance (Fisher’s Exact Test, P = 0.58), while HAQERs were significantly more likely to be differentially active than expected by chance (Fisher’s Exact Test, Odd’s ratio 2.88, P < 0.01).

**Figure 4.**
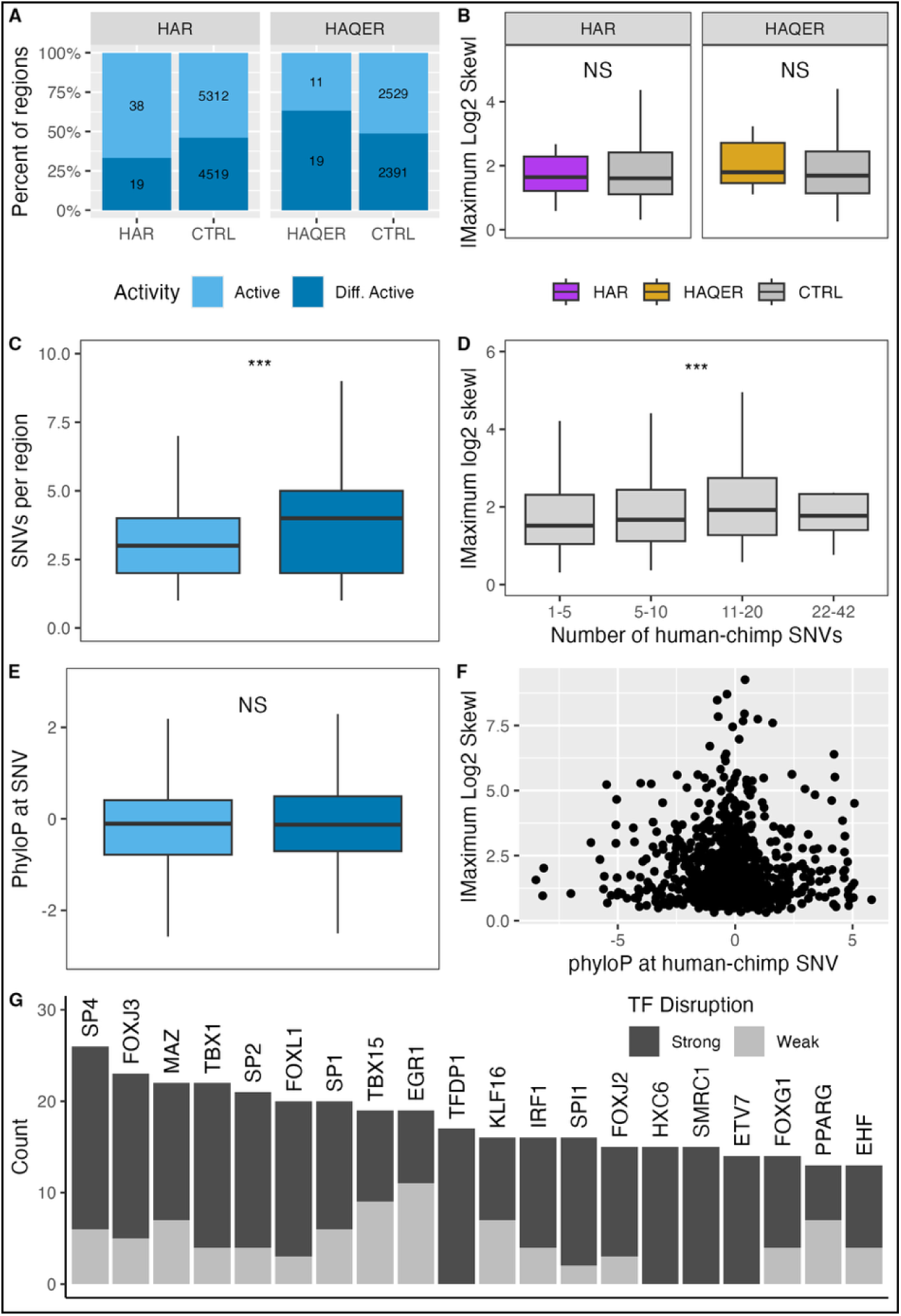
Association of sequence features with differential activity. (A) rates of differential activity in HAR/HAQER overlapping regions and matched control sets (CTRL). (B) distribution o maximum magnitude of the log2 skew across cell types for differential activity regions overlapping a HAR or HAQER and matched control sets. (C) number of human-chimp SNVs in active or differentially active directly mappable tiles, *** P < 5 x 10^15^, Wilcoxon rank-sum test. (D) boxplots of the magnitude of the log2 skew for directly mappable tiles containing different numbers of SNVs, * = P < 0.01, Pearson’s correlation. (E) boxplots of phyloP scores for SNV positions in tiles with only a single SNV. (F) relationship between phyloP scores and magnitude of the maximum skew across cell types for SNV positions in tiles with only a single SNV. (G) number of strongly and weakly disrupted TF binding motifs in differentially active tiles with only a single SNV. NS = not significant. Outliers for B-E are not shown.

Both HARs and HAQERs are defined based on specific genomic properties and characterized by numerous sequence differences. Therefore, to evaluate these methods’ ability to predict differential activity, we matched our HAR/HAQER overlapping regions with control sets comprising other tested regions from our MPRA with similar sequence features (see Methods). Compared to their respective control sets, neither regions overlapping HARs nor HAQERs were more likely to be differentially expressed (Fisher’s exact test, P > 0.05, Figure 4A). To compare the magnitude of differential activity associated with HAR and HAQER overlaps, the absolute value of the maximum log2 fold skew across either tile in all cell types was used. Once again, neither regions overlapping HARs nor HAQERs showed significantly different maximum magnitudes of differential expression compared to their control set (Wilcoxon rank-sum test, P > 0.1, Figure 4B).

As HARs and HAQERs explain only a small fraction of the differentially active tiles, we further explored the sequence features driving differential expression. We first looked at tiles directly mappable across species (contiguous, orthologous sequences of equal length). On average, differentially active tiles contained significantly more SNVs than all active sequences (Wilcoxon rank-sum test, P < 5 x 10^15^, Figure 4C). Both the sets of differentially active and active sequences had significantly more SNVs on average than the total set of tested sequences (Wilcoxon rank-sum test, P < 5 x 10^15^). Further supporting an association between the number of SNVs and differential expression, we found a significant correlation between sequence divergence and the maximum magnitude of change in enhancer activity observed in any cell type with a consistent skew direction (Pearson’s R = 0.033, P < 0.01, Figure 4D), in agreement with a previous study (Klein et al. 2018). As HAQERs are some of the fastest evolving regions in the human genome, the pattern of differentially activity described above is consistent with the large number of SNVs distinguishing the human and chimp sequences at these regions.

### Features of single variant tiles

While most of the tested tiles had multiple differences between the human and chimp sequence (84.4%), either due to SNVs or small indels, 20,335 orthologous tile pairs were distinguished by only a single SNV (15.6%). As this set contains only a single locus which can explain any differential activity, we analyzed the 1,030 differentially active tiles with only a single SNV in greater detail. First, we analyzed phyloP as phylogenetic conservation scores are often used to identify regulatory elements or critical base pairs therein. For all active tiles with just a single SNV, we found that phyloP was not significantly associated with activity (Figure 4E, Wilcoxon rank-sum test, P = 0.36). Within the set of differentially active tiles, we tested for correlations between the phyloP score and maximum magnitude of change in enhancer activity but found none (Figure 4F, Pearson’s correlation, P = 0.95). This result is unsurprising given the close relationship between humans and chimps that makes the two species likely to share many lineage-specific regulatory elements with low phyloP scores. We next determined which TFs have predicted binding disruptions in sequences containing only one SNV. Overall, we found that our set of tiles containing a single SNV showed strong disruptions of predicted TF motifs for 511 TFs and weak disruption of 679 TFs (Tables S10-S11). The top three most frequently disrupted strong TFs and weak TFs were *SP4, FOXJ3, TBX1* and *EGR1, PURA, TBX15,* respectively (Figure 4G). *TBX1*, *EGR1,* and *TBX15* have previously been shown to have important roles in skeletal development (Singh et al. 2005; Reumann et al. 2011; Funato et al. 2015). While not one of the most frequently disrupted TF motifs, *CTCF* binding was disrupted strongly four times and weakly twice, consistent with our results above. The disruption of binding sites for these skeletal development transcription factors provides a potential molecular mechanism by which single nucleotide changes could alter skeletal gene expression patterns between humans and chimpanzees.

## Discussion

Using the MPRA, we comprehensively tested 70,000 genomic regions accessible across the developing human postcranial skeleton for differential activity between humans and chimpanzees. By testing our library in three skeletally relevant cell lines, two chondrocyte lines and one bone marrow derived leukocyte line, we found that >30,000 regions (45% of the total) showed enhancer activity. As the library was designed using genomic regions accessible in developing cartilage, inactive sequences may not be functional in the tested MPRA cell types due to differences in the cellular environment or have a repressive function, something our MPRA design is insufficiently powered to detect due to the low activity of the basal promoter (Tewhey et al. 2016). We see no strong evidence of bias towards activity of human regions compared to chimp, consistent with previous studies that have compared activity in both human and chimp cell lines (Whalen et al. 2023) or used mouse cells as a neutral trans environment (Girskis et al. 2021).

Of the active regions, we found that 11,542 (37.6%; or 17% of the entire pool) were differentially active between human and chimpanzee, potentially contributing to skeletal differences between the species. This extensive functional divergence suggests that the evolution of human-specific skeletal features involves widespread regulatory changes rather than a few modifications to key developmental pathways. This is consistent with recent work by Senevirathne et al. (Senevirathne et al. 2025) on the pelvic ilium showing overlaps of key regulatory elements with many sequence differences. Even though our library tested regions with known accessibility in particular bones of the developing human skeleton, we did not observe clear patterns across skeletal elements. Differentially active regions in each cell type had a similar distribution of overlaps with skeletal tissue accessibility to that present in the entire tested library. This suggests that none of the tested cell types closely matches any of the developmental tissues. In support of this, we found that regions identified at the later stage were more likely to show activity (Figure 2H). As these regions are identified later in development, they should reflect greater cell differentiation and thus better match the tested cell lines, all of which were derived from tissues collected at much later stages (range 18 weeks gestation to 53 years old). A substantial degree of timepoint specificity in activity (and therefore differential activity) is expected, as a study of activity changes in human brain organoids using lentiMPRA found that 75% of the tested enhancers had activity only at a specific time point (Capauto et al. 2024). While the skeletal system may have relatively few cell types, the need to precisely shape and grow individual bones and joints requires exact spatial and temporal regulation across the body to ensure proper function (Guenther et al. 2008; Richard et al. 2020; Muthuirulan et al. 2021; Coveney et al. 2025). Future experiments aiming to test the inactive sequences should prioritize cell lines with expression of the TFs that bind motifs that are present in the library but depleted in the active sequences in this experiment such as *NFIC, NFIA, NFIX,* all of which play important roles in the regulation of early skeletal development (Driller et al. 2007; Singh et al. 2018; Lee et al. 2020; Kooblall et al. 2023). Our finding that activity and differential activity were correlated with the number of oligos tested for each skeletal element may also be a product of our MPRA approach. Our library was tested in an episomal context rather than via genomic integration, as in the lentiMPRA method, which has been shown to correspond better to functional genomic signals (Inoue et al. 2017). As our library was entirely based on ATAC-seq data, the lack of chromosomal integration may have reduced the power of our assay to test important subsets of the regions that are functional *in vivo*.

Despite these limitations, we nonetheless identified over 11,000 regulatory elements with sequence differences between humans and chimpanzees that lead to differential regulatory activity. These elements are accessible across the postcranial skeleton, with some accessible in all tissues and others in just one. This is certainly an underestimate of the total number of active enhancers for the reasons stated above as well as the size of test constructs. While we tested a centered 540 base pair window for each element, our split 270 tile design may split active elements into inactive parts or fail to capture longer functional sequences (Girskis et al. 2021).

While directly tying these differentially active regions to their impacts on skeletal morphology and evolution remains a major challenge, this result is consistent with a complex, polygenic basis for human skeletal evolution (Okamoto et al. 2025; Senevirathne et al. 2025; Richard et al. 2025). Future experimental work is needed to understand the phenotypic implications of each of these regulatory changes.

In comprehensively testing skeletal regulatory elements between human and chimpanzee, we also sought to compare the efficacy of commonly used methods for identifying regions likely involved in the evolution of human-specific traits (Pollard et al. 2006; Prabhakar et al. 2006; Bird et al. 2007; Bush and Lahn 2008; Gittelman et al. 2015; Kostka et al. 2018; Mangan et al. 2022; Bi et al. 2023; Whalen et al. 2023; Keough et al. 2023; Yoo et al. 2025). We found that active tiles containing a HAR were not more likely to be differentially active than expected by chance, and that differentially active regions overlapping HARs have similar skew sizes to matched differentially active regions. While some studies prioritizing HARs have identified key genomic changes underlying human-specific phenotypes (e.g., (Aldea et al. 2021)), the overall success rate of tested HARs is poor (Baumgartner et al. 2024). While HARs are identified due to an excess of mutations, these mutations do not necessarily have synergistic effects and in many cases have been shown to compensate for one another, leading to multiple changes that individually alter gene expression, but have little effect when considered together (Whalen and Pollard 2022; Whalen et al. 2023). In contrast, we found that HAQERs significantly predicted differential activity but again did not show exaggerated skews. As HAQERs are the fastest evolving regions of the human genome and therefore contain many sequence differences, this finding is consistent with the overall higher number of SNVs observed in differentially active sequences. Our study design prevents interrogation of the magnitude and direction of effect of each SNVs within the same tile so an important future direction for interrogating these regions would be a saturation mutagenesis assay to explore the individual and additive effects of neighboring variants (Klein et al. 2018; Siraj et al. 2024). Our finding that HARs poorly predict human-specific activity and that the total number of differences is a stronger predictor may be due the specific assay we performed or be limited to the skeletal system and not apply to other tissues such as the brain. Future studies using similarly comprehensive enhancer screens in other tissues are needed to understand how the relationship between sequence changes and enhancer activity levels differs by anatomical system.

In summary, this study identified thousands of regulatory elements involved in early skeletal development with differential activity between humans and chimpanzees, providing a rich resource for understanding human adaptations across the postcranial skeleton as well as for comparing the efficacy of computational methods for identifying genomic regions of particular relevance to human-specific biology. Our results highlight the challenge of predicting functionally important changes from genomic sequences alone as even a few base pair changes can have large effects on enhancer activity.

## Methods

### MPRA library design

Putative regulatory elements identified by ATAC-seq on cartilage tissue at two developmental timepoints, the first at ∼50-60 embryonic days and the second at ∼70 embryonic days, were merged to generate a single composite dataset for the developing human postcranial skeleton. Skeletal elements included the ATAC-seq data from the phalanges and metapodials of the hands and feet (Okamoto et al. 2025), proximal and distal ends of the humerus, femur, ulna, and tibia (Richard et al. 2020), the ilium, ischium, pubis, and acetabulum (Young et al. 2022), the head, neck, and blade of the scapula, as well as thoracic and lumbar vertebrae (second timepoint only) (Richard et al. 2025). This set of genomic regions was filtered using BEDTools (Quinlan and Hall 2010) to remove regions that share accessibility with embryonic brain tissue. Each putative regulatory element was then standardized to a length of 540 base pairs (bps) and divided into two 270 bp tiles meeting at the center of the region. The genomic sequence for each tile was extracted from the hg38 human reference genome using the *getSeq* function from the *R Biostrings* package (version 2.68.1) (Pagès et al. 2023). To get the corresponding chimpanzee genomic sequence for each tile, the coordinates were lifted over to the chimpanzee genome (PanTro6) and the chimp sequence extracted as above. Any tiles that failed to liftover to chimp or had identical sequences between the two species were removed. Additionally, any tiles that mapped to regions on multiple chimp chromosomes were removed since the sequence is not contiguous (possibly due to a chromosomal rearrangement at this position) and therefore any difference in expression between the two species would be difficult to interpret biologically. This approach of lifting over human regions produced a human-bias in the library, such that chimpanzee specific-sequences that do not align to human (i.e., human deletions) were not included in the oligos. To avoid comparing sequences of substantially different lengths, only regions with a chimp sequence at least 90% the length of that seen in humans were included.

15-bp adapters were then added to both ends of each sequence: 5′-ACTGGCCGCTTGACG and CACTGCGGCTCCTGC-3′. If the chimp sequence was shorter than the human, the chimp sequence was padded with additional vector sequence to ensure a uniform size for each oligo of 270 bps. 75 positive and 295 negative controls provided by Ryan Tewhey were added to the oligonucleotide pool. The final pool of 290,000 oligos was synthesized by TwistBioscience (Table S12).

### MPRA library construction

Library construction followed an established protocol from the Tewhey lab, MPRAv3 Library Generation Protocol (v06/2021), detailed in (Siraj et al. 2024). Briefly, unique 20 bp barcodes were added to the oligos using PCR (15 cycles following (Xue et al. 2023)) before insertion into SFiI digested pMPRAv3:Δluc:ΔxbaI vector by Gibson assembly. After determining the colony-forming unit (CFU) count necessary to achieve a library composition of a least 50 barcodes per oligo (Lagunas et al. 2023; Delfosse et al. 2023), we performed five electroporations of 1μl ligated vector into 50 μL 10-beta electrocompetent cells (NEB) before recovery, 16 hours of growth at 30°C in six 500 mL flasks of TB media with 100 ug/ml carbenicillin, and purification using a Qiagen Plasmid Plus Giga Kit. Library composition and diversity were confirmed by CFU plates and colony PCR followed by Sanger sequencing. Oligo sequences were associated with barcodes by sequencing the plasmid library on an Illumina Novaseq X 25B flowcell to a depth of 978.5 million reads.

To construct the final MPRA library, a GFP amplicon containing a minimal TATA promoter (amplified from pMPRAv3:minP-GFP, Addgene #109035) was inserted into 10ng of AsiSI-digested plasmid library. After SPRI cleanup, the library was transfected into 10-beta competent cells, split into six 2mL cultures and recovered at 37°C for one hour followed by expansion of each culture in 500 mL of TB media with 100ug/ml carbenicillin. After 16 hours of growth at 30°C, plasmid was purified using ZymoPURE II Plasmid Gigaprep kits and stored at - 20°C for use in transfections.

### MPRA library transfections

Three cell-types were used for MPRA transfections, two chondrocyte lines (T/C-28a2 and CHON-002) and a lymphoblast line (K562). 300-500 million cells per technical replicate were transfected at 200ug/ml (K562, T/C-28a2) and 300ug/ml (CHON-002) of plasmid library, using a MaxCyte ATx electroporation device. Preexisting electroporation protocols established by MaxCyte were used and optimized in our hands by FACS analysis using a control plasmid on the device for each cell line. R1000 cartridges were used at ∼125 million cells per cartridge. Following electroporation, cells were allowed to rest for 25 minutes in proprietary MaxCyte Buffer in a 6-well plate at 37°C. Cells were then plated in their respective media at a 50%-70% density. Cells were harvested 24 hours post-transfection, spun down and snap frozen on dry ice before storing at −80°C. Five technical replicates were transfected per cell line.

### RNA extraction

Pellets were defrosted before the addition of RLT buffer containing 2M DTT (40ul for every 1ml of RLT buffer for >2×10^8 cells). Pellets were disrupted using a 10 mL syringe and 21-gauge needle before the addition of equal amounts of 70% ethanol. Samples were then processed using the Qiagen Maxi RNeasy and following the isolation procedure instructions. RNA was nano dropped and stored at −80°C.

### Isolation of GFP containing RNA

Half of the RNA sample was taken for MPRA clean up processing and library generation whilst saving the remaining half. Samples underwent a 1-hour DNase digest at 37°C to remove any contaminating DNA, particularly coming from the plasmid. Following this biotagged *GFP* primer probes (CGCCGTAGGTGAAGGTGGTCACGAGGGTGGGCCAG/3BioTEG/, CCAGGATGTTGCCGTCCTCCTTGAAGTCGATGCCC/3BioTEG/, CCTCGATGTTGTGGCGGGTCTTGAAGTTCACCTTG/3BioTEG/) were adhered to the *GFP* sequence of mRNA products using formamide cross linking (2.5 hours at 65°C). This was followed by hybridization of streptavidin magnetic Dynabeads^TM^ (15 mins, rtp, on nutator) that bind to the BioTEG portion of the probes now bound to the mRNA products of interest.

Dynabeads were made RNase free prior to use following provided protocols. mRNA bound to Dynabeads was then isolated using a magnet to remove beads from the rest of the RNA containing solution in clean up stages followed by elution in 25ul of RNAse free water at both 70°C and then 80°C.

To remove any remaining plasmid DNA, a further 1-hour 37°C digest was performed before RNA SPRI bead cleanup (standard isolation and elution protocols) and performing first strand synthesis and reverse transcription (ThermoFisher Scientific, 18080044). Following reverse transcription, DNA SPRI was performed to remove any remaining reagents and concentrations checked using qPCR analysis. If CT value was below 18, samples were amplified at 2 cycles above their CT value before attaching unique illumina P5 and P7 adapter pairs. At this stage, four plasmid replicates were included and prepared for sequencing to assess library diversity representation for downstream processing. Samples were then assessed for concentration before pooling to form an equimolar pool. The resulting pool was sequenced five times on an Illumina NovaSeq X 10B flowcell to a depth of >125 million reads per replicate.

### MPRA data processing

Data processing was performed using computational pipelines developed by the Tewhey lab for MPRA analysis (https://github.com/tewhey-lab/MPRASuite/tree/main) (Tewhey et al. 2016; Siraj et al. 2024). Briefly, unique oligo:barcode pairs were identified and used to generate count tables for each oligo in each RNA and plasmid replicate. The RNA:plasmid ratios were then statistically compared to identify active and differentially active sequences. For the first step, oligo sequences were associated with barcodes using the MPRAmatch pipeline with the following modifications. Since the Novaseq X only generates reads of 150 bp pairs in length, the paired end reads did not overlap and did not cover the entire barcoded oligo sequence (total length: 369 bp, oligo (270bp) + F primer (22bp addition) + R primer (77bp addition)). Therefore, we modified the MPRAmatch script, removing the initial step that calls FLASH2 to merge overlapping pair end reads, instead merging the forward and reverse reads with an intervening 100 Ns using the BBMap fuse.sh tool. To avoid an error due to the incomplete coverage of the oligo sequences, the standard MPRAmatch “%sequence mismatch” threshold was circumvented by setting the value for all sequences below the standard threshold. This approach merged the oligo sequence to the barcode information and allowed for unique barcode identification for most oligos. As this study compares human and chimp genomic differences, many tiles had multiple differences rather than just a single SNP, enhancing our power to distinguish between the human and chimp oligos for the same tile. While a small number of sequences with unique features only at the end of the tile were lost, this approach allowed us to take advantage of the high number of reads enabled by the Novaseq X platform for identifying barcodes attached to longer oligos, which are more biologically informative than shorter sequences (Klein et al. 2020).

After sequencing of the RNA and plasmid libraries, paired end reads were merged using NGmerge (Gaspar 2018). Read counts for each barcode were generated using the MPRAcount pipeline. Differential expression between the human and chimp versions of each oligo was determined using the MPRAmodel pipeline in R (version 4.4.1). This pipeline uses DESeq2 for normalization and statistical analysis using RNA/DNA ratios for each oligo and a negative binomial generalized linear model (GLM) to identify changes in activity levels. Tiles were defined as active if they had a |log2[fold-change]| > 1 and Bonferroni-adjusted p-value < 0.01, and differentially active if either the human or chimp tile was active and the Benjamini-Hochberg-adjusted p-value for a non-zero species effect was < 0.1. For some regions, despite lifting over successfully (i.e., mapped to a sequence of similar length) and passing other quality filters, alignment quality was very poor between the human and chimp sequences as calculated by the Biostrings pairwiseAlignment function in R (Pagès et al. 2023). Since these sequence differences are potentially due to problematic genome alignments rather than regulatory evolution, any active or differentially active sequences with an alignment score ≤ 0 were removed (14,284 pairs, 9.9% of all tested pairs). Statistical test results for activity and differential activity of all tiles for each cell line are available in Tables S13-S18.

### Transcription factor (TF) binding motif enrichment analysis

TF binding motif enrichments for region sets were calculated in R using MonaLisa (Machlab et al. 2022) with human PWM matrices from the 2020 JASPAR database (Fornes et al. 2019). Statistical significance was assessed using a negative log10 cut-off of 4. To determine which TF motifs were differentially bound due to SNVs in tiles with only a single human-chimp SNV, we used the *motifbreakR* (version 2.14.2) *snps.from.rsid* function (Coetzee et al. 2015) with *BSgenome.Hsapiens.UCSC.hg38* to get the human sequence surrounding each SNV. Motif enrichments for each SNV were identified using the *motifbreakR* function with the following parameters: filterp = TRUE, threshold = 1e-4, method = “ic”, bkg = c(A=0.25, C=0.25, G=0.25, T=0.25), BPPARAM = BiocParallel::bpparam()).

### Tests for enrichments

Human accelerated regions (HARs) were compiled from refs. (Pollard et al. 2006; Prabhakar et al. 2006; Bird et al. 2007; Bush and Lahn 2008; Gittelman et al. 2015; Kostka et al. 2018; Bi et al. 2023). Human ancestor quickly evolved regions (HAQERs), the fastest evolving regions of the human genome, were from ref. (Yoo et al. 2025). The HAQERs were lifted over to hg38 using the liftover file available at https://github.com/marbl/CHM13. Significance of enrichment were identified using Fisher’s exact test via the *binom.test* function in R. Significance of distribution shifts were identified using a two-tailed Wilcoxon rank-sum test via the *wilcox.test* function in R. Gene ontology analysis of genomic region sets was performed using GREAT (v4.0.4), which identifies potential target genes for genomic regions and tests their enrichment for biological ontology annotations, including human and mouse phenotypes (McLean et al. 2010). In general, active sets were enriched against the total tested library and differentially active sets were tested against the appropriate active set to control for bias in the sequences tested.

Testing for enrichments in the HARs and HAQERs requires special consideration of the appropriate background. To generate a matched control set for differentially active tiles overlapping a HAR, active tiles had to not overlap a HAR and have only 5-7 base pair differences or a deletion of chimp sequence less than 5 base pairs, reflecting the profile of the HAR overlapping tiles. To generate a matched control set for differentially active tiles overlapping a HAQER, active tiles had to not overlap a HAQER and have at least 8 base pair differences or a deletion of chimp sequence larger than 1 base pair, reflecting the profile of the HAQER overlapping tiles.

### Measures of phylogenetic conservation

PhyloP scores for regions were calculated as the mean across a tile using the Zoonomia phyloP scores (Sullivan et al. 2023) and shifts in phyloP between sets of tiles were calculated using two-sided Wilcoxon rank-sum tests. PhyloP scores for single positions were extracted from the same resource.

### Proximity to TSS locations

The minimum distance of each region to the nearest TSS as defined in the refTSS database (Abugessaisa et al. 2019) was calculated using *bedtools closest -d*. Differences in TSS distances were compared between sets using a two-sided Wilcoxon rank-sum test.

### Data access

All raw and processed sequencing data generated in this study have been submitted to the NCBI Gene Expression Omnibus (GEO; https://www.ncbi.nlm.nih.gov/geo/) under accession number GSE298093. The code used to perform these analyses and to generate these figures is available at: https://github.com/aokamoto-bio/ human_skeletal_evolution_MPRA.

## Acknowledgments

We would like to thank Ryan Tewhey, James Xue, Daniel Lieberman, Kimberly Cooper, Clifford Tabin, David Gokhman, and members of the Capellini lab for critical insight and commentary on this project. A.S.O and T.D.C. were supported by a National Science Foundation Doctoral Dissertation Improvement Grant (BSC-2337516). A.S.O. was supported by a National Science Foundation Graduate Research Fellowship (DGE-1745303). T.D.C. was supported by the American School of Paleontological Research (ASPR) via Harvard University.

## Author Contributions

A.S.O. designed and constructed the MPRA library, assisted with transfections and RNA isolations, performed all computational analyses, contributed to study design, secured funding for the project, wrote the manuscript, and generated figures with input from all authors. C.R.C performed all transfections and RNA isolations. D.S.G. aided in transfections and RNA isolations. T.D.C. conceived of and supervised all aspects of the project, designed the study, secured funding for the project, and edited and revised the manuscript with input from all authors.

