## Supplemental Text and Figures for "Massively parallel functional screen identifies thousands of regulatory differences in human versus chimpanzee postcranial skeletal development"

**Species-specific conserved element deletions**

The library design used in this experiment was biased to the human sequence and therefore genomic regions lost in humans were not included in the corresponding chimpanzee tiles (due to direct sequence mapping rather than mapping region endpoints during liftover). Chimpanzee losses were included, however. These nucleotides may be ‘lost’ due to human gains, chimp losses, or alignment issues. When analyzing tiles with a chimp deletion, there was no significant difference in the size of the deletions associated with activity or differential activity (P > 0.05, Wilcoxon rank-sum test, Figure S7A). Similarly, there was no correlation between the size of the deletion and the magnitude of differential activity (P > 0.2, Pearson’s correlation, Figure S7B).

Some species-specific deletions occur in regions that are otherwise highly conserved in other vertebrates, such as human conserved element deletion (hCONDEL) events (McLean et al. 2011; Xue et al. 2023). While these elements were not included in our library, preventing direct analysis of hCONDELs and comparisons of efficacy with HARs and HAQERS, our library design strategy could potentially include chimpanzee conserved element deletion events (cCONDELs). As a corresponding set of cCONDELs to the hCONDELs from (Xue et al. 2023) are not available, we instead investigated a comparable set of tiles within our MPRA as those with a loss of alignable sequence in chimp and a positive phyloP score averaged across the tile (as 266/298 (89%) of the regions that span an hCONDEL position had positive phyloP scores). Altogether 2,150 active tiles met these criteria, of which 473 exhibited differential activity (28.2%). This is comparable to the 33% of active HARs with differential activity. There was no significant difference in the size of the deletions associated with activity or differential activity (P > 0.2, Wilcoxon rank-sum test, Figure S7C) or correlation between the size of the deletion and the magnitude of differential activity (P > 0.7, Pearson’s correlation, Figure S7D). While our library was not designed to test deleted sequences and there may be some effects of the padding added to the edges of oligos containing chimp deletions to ensure constant oligo size or nearly SNVs also present in the oligo, deletions are not particularly associated with differential activity in this dataset.

**Supplemental Figures**

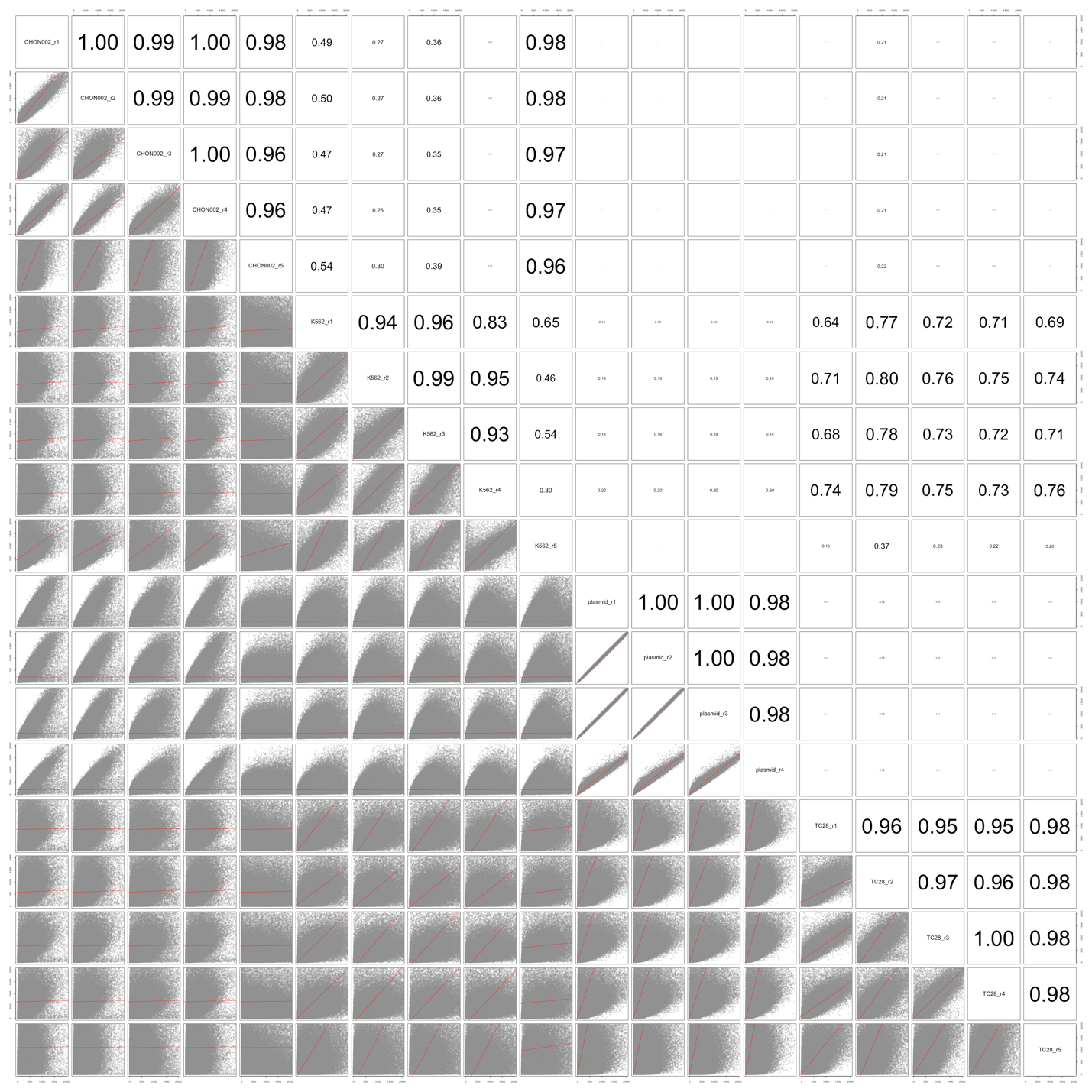

**Figure S1.** Correlation of oligo counts for each replicate. Pearson’s R shown for each pairwise comparison.

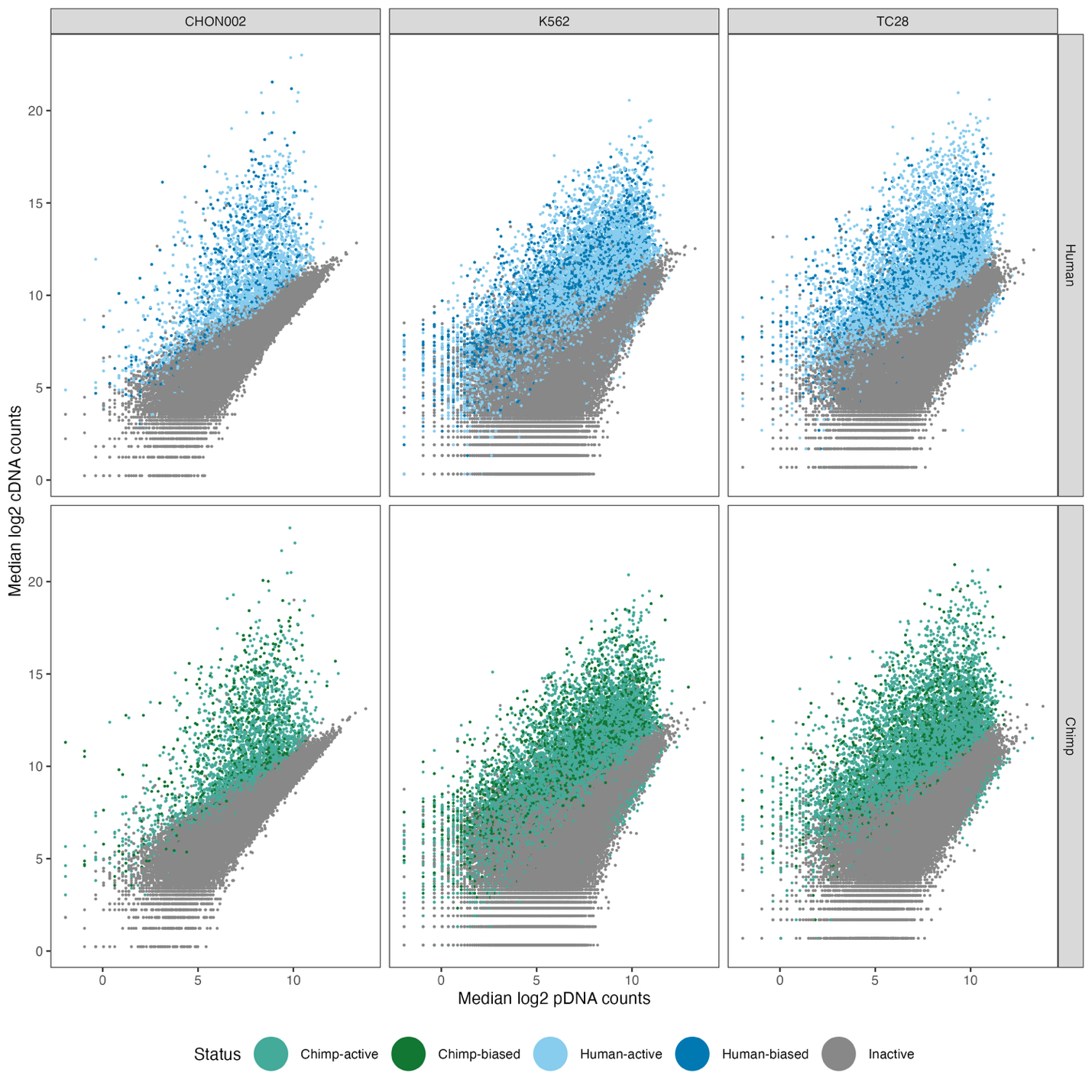

**Figure S2.** Quantification of enhancer activity via MPRA. Plasmid DNA and cDNA counts for tested human and chimp sequences (top and bottom, respectively) in each cell type showing the number of sequences with activity (lighter color) or differential activity between species (darker color).

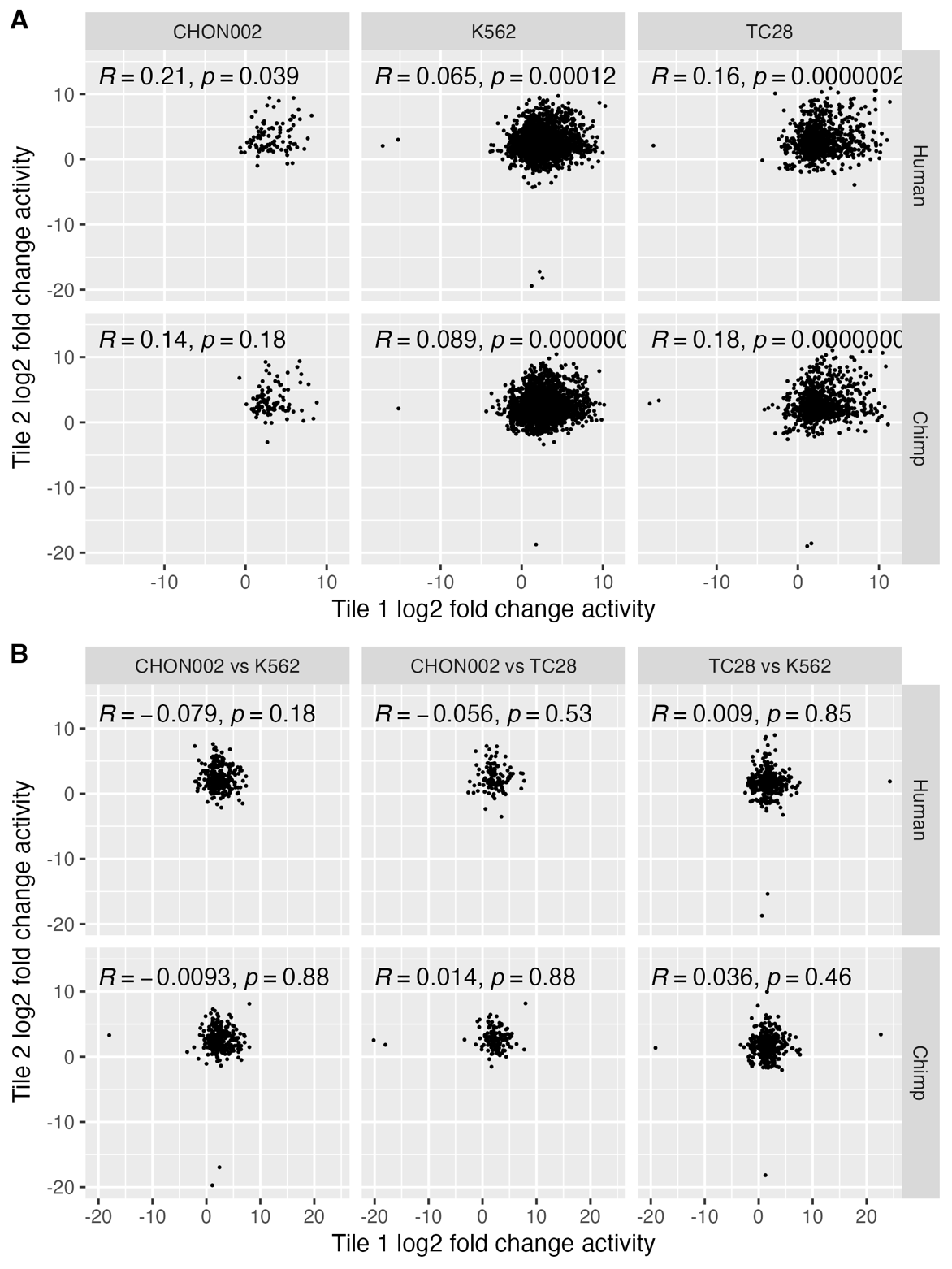

**Figure S3.** Correlation of activity between tiles within regulatory regions. (A) correlation of activity of adjacent tiles with activity within a single cell line, shown separately for both the human and chimpanzee versions of each sequence. (B) correlation of activity of adjacent tiles that were active in different cell lines, shown separately for both the human and chimpanzee versions of each sequence.

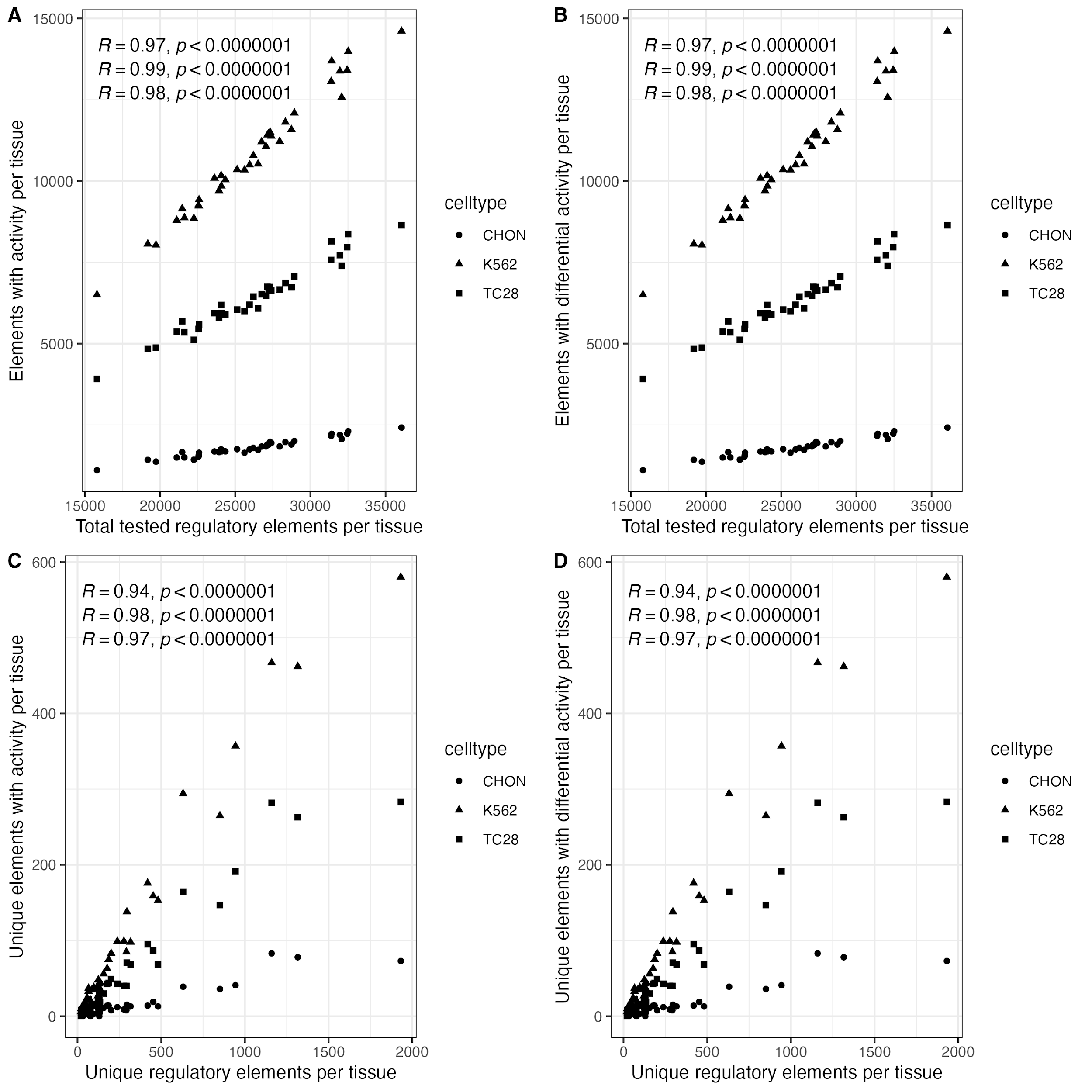

**Figure S4.** Correlation of regulatory element size and MPRA results by cell type. (A) correlation of elements with activity or (B) differential activity in each cell type per tissue with the total number of tested regulatory elements for that tissue. (C) correlation of tissue-unique elements with activity or (D) differential activity in each cell type with the total number of tested unique regulatory elements for that tissue. Statistics for Pearson’s correlation shown.

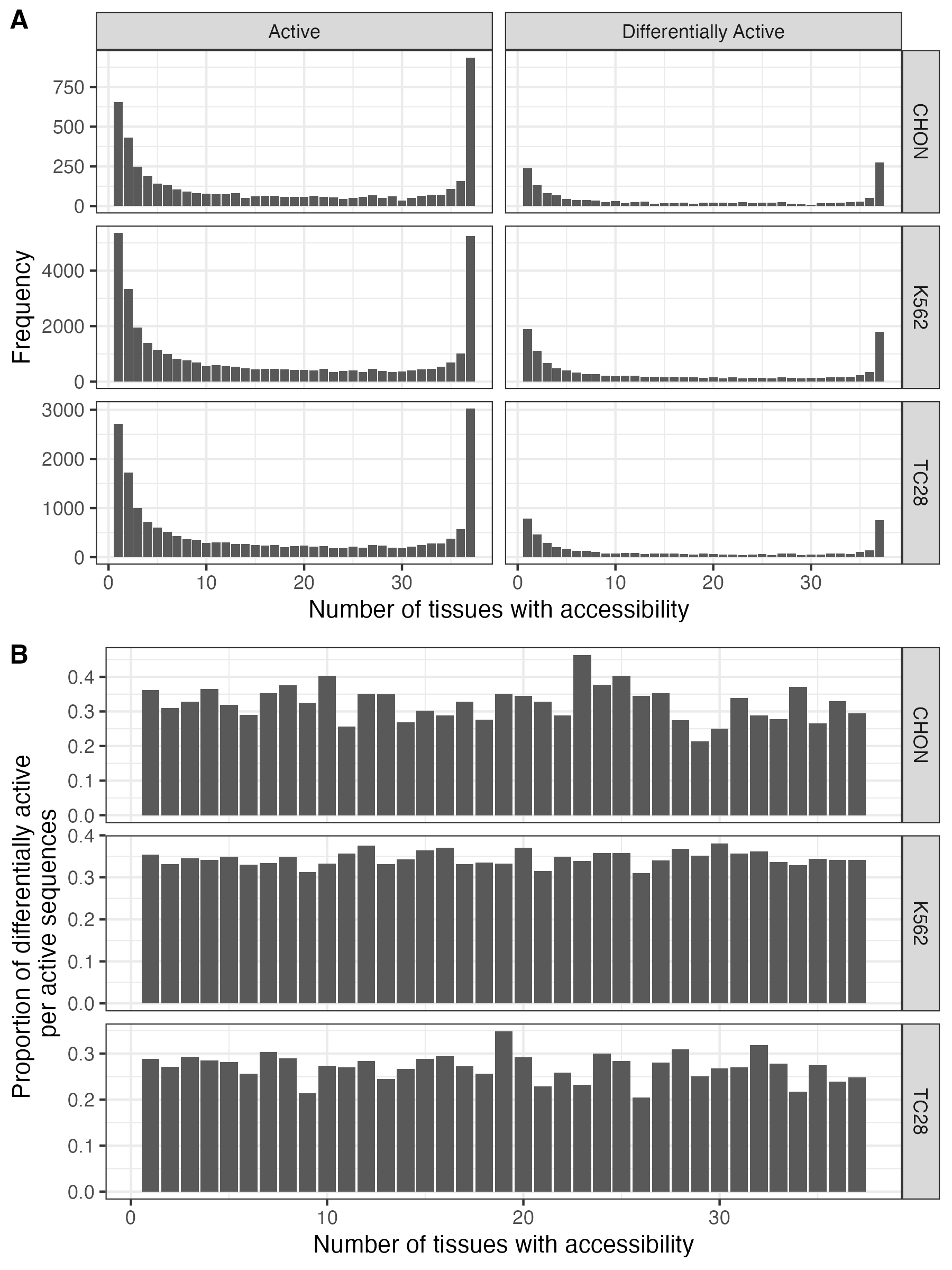

**Figure S5.** Accessibility sharing of active and differentially active regions. (A) Frequence of active/differentially active elements falling in regions with different levels of shared accessibility. (B) proportion of active regions that are differentially active for each cell type.

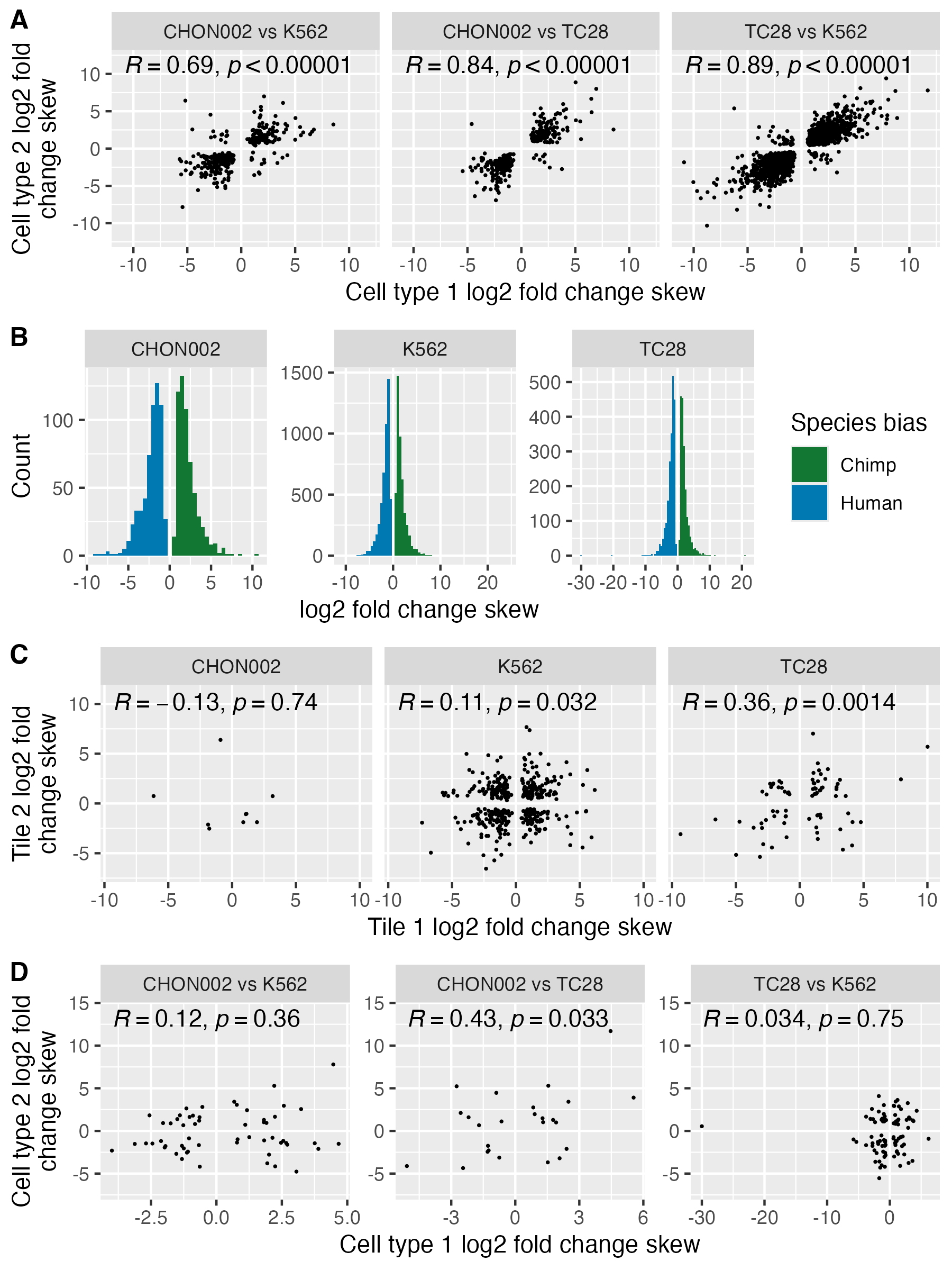

**Figure S6.** Correlation of differential activity across cell lines. (A) correlation of skew for tiles with differential activity in multiple cell lines. (B) distribution of skews in differentially active tiles for each cell line. (A) correlation of skew for tiles with differential activity in multiple cell lines. (C) correlation of skew of adjacent tiles with differential activity within a single cell line. (D) correlation of skew of adjacent tiles that were differentially active in different cell lines. Statistics for the Pearson’ correlation shown in A,C,D.

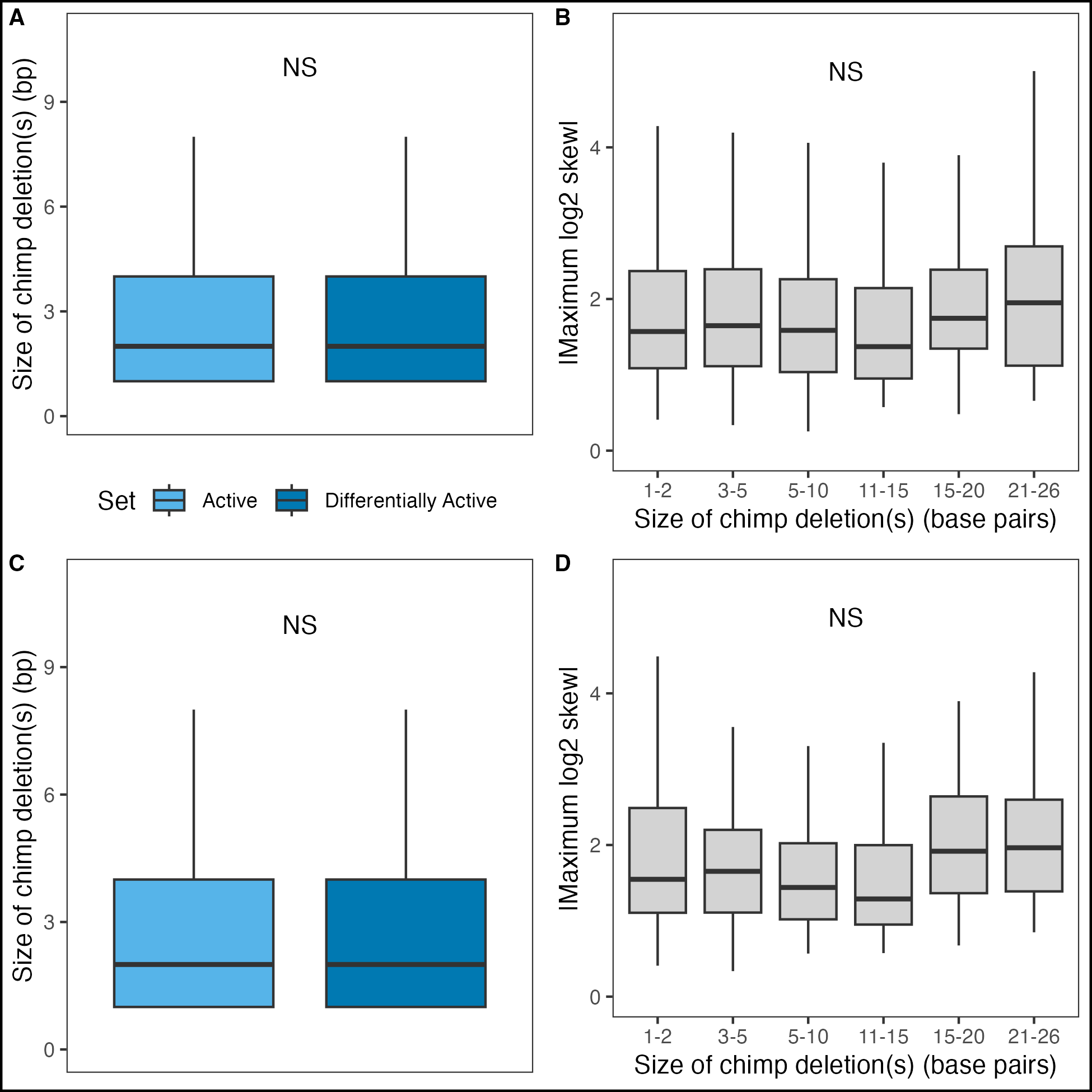
**Figure S7.** Features of tiles with chimp sequence deletions. (A) size of chimp sequence deletions in active or differentially active tiles. (B) boxplots of the magnitude of the log2 skew for tiles containing different lengths of chimp deletions. (C) size of chimp sequence deletions in active or differentially active tiles with positive average phyloP scores. (D) boxplots of the magnitude of the log2 skew for tiles containing different lengths of chimp deletions with positive average phyloP scores.
